# *Escherichia coli* YigI is a conserved ɣ-proteobacterial acyl-CoA thioesterase permitting metabolism of unusual fatty acid substrates

**DOI:** 10.1101/2022.01.10.475765

**Authors:** Michael Schmidt, Theresa Proctor, Rucheng Diao, Peter L. Freddolino

## Abstract

Thioesterases play a critical role in metabolism, membrane biosynthesis, and overall homeostasis for all domains of life. In this present study, we characterize a putative thioesterase from *Escherichia coli* MG1655 and define its role as a cytosolic enzyme. Building on structure-guided functional predictions, we show that YigI is a medium- to -long chain acyl-CoA thioesterase that is involved in the degradation of conjugated linoleic acid (CLA) *in vivo*, showing overlapping specificity with two previously defined *E. coli* thioesterases TesB and FadM. We then bioinformatically identify the regulatory relationships that induce YigI expression, which include: an acidic environment, high oxygen availability, and exposure to aminoglycosides. Our findings define a role for YigI and shed light on why the *E. coli* genome harbors numerous thioesterases with closely related functions.

**Importance:** Previous research has shown that long chain acyl-CoA thioesterases are needed for *E. coli* to grow in the presence of carbon sources such as conjugated linoleic acid, but that *E. coli* must possess at least one such enzyme that had not previously been characterized. Building off of structure-guided function predictions, we showed that the poorly annotated protein YigI is indeed the previously unidentified third acyl CoA thioesterase. We found that the three potentially overlapping acyl-CoA thioesterases appear to be induced by non-overlapping conditions, and use that information as a starting point for identifying the precise functions catalyzed by each such thioesterase, which is an important prerequisite for their industrial application and for more accurate metabolic modeling of *E. coli*.

## Introduction

Thioesterases play an important role in fatty acid metabolism, with thioesterase activity involved in the release of newly synthesized molecules from an acyl carrier protein [1] in many organisms. Other thioesterases are often involved in maintaining homeostasis of free and activated fatty acids [2] and salvaging acyl-CoA intermediates of fatty acid breakdown that cannot be subjected to additional rounds of beta oxidation (e.g., [3]). While the commonly studied bacterium *Escherichia coli* does not require a thioesterase to release fatty acids from the acyl carrier protein, instead relying on direct trans-esterification of the new fatty acid to other targets such as glycerol 3-phosphate [4,5], it is still dependent upon several thioesterases to carry out other essential metabolic reactions. The *E. coli* K12 MG1655 genome encodes several characterized thioesterases, including the periplasmic multifunctional TesA [6,7], the cytosolic TesB (Thioesterase II) which acts on medium- to long-chain acyl CoA substrates [8], the cytosolic FadM (Thioesterase III) that prefers long-chain derivatives [9,10], and the cytosolic YciA that harbors broad substrate specificity [11]. MG1655 also encodes several specialized thioesterases known to act along specific metabolic pathways, including MenI which is important for menaquinone synthesis [12], PaaY and PaaI which are both involved in phenylacetate catabolism [13], and EntH which plays an important role in enterobactin synthesis [14]. Importantly, individual knockouts of the broad-specificity cytosolic *E. coli* thioesterases (TesB, FadM, or YciA) are not significantly detrimental to the organism, implying that there is some pathway compensation across this enzyme family [9,11,15].

The acyl CoA thioesterases TesB and FadM were previously shown to enable the cell to recycle CoA from intermediates of fatty acid beta oxidation that cannot be further processed through that pathway [3,10]; at the same time, genetic studies indicated that at least one more thioesterase with overlapping substrate specificity must exist, given the ability of even a *tesB fadM* double mutant to grow on conjugated linoleic acid (CLA) [3]. In a recent whole-proteome application of the COFACTOR2 workflow for combined structure prediction and functional annotation to *E. coli* MG1655, we identified the poorly annotated gene *yigI* as a likely acyl CoA thioesterase, thus indicating that it is a candidate for the additional thioesterase previously posited in Nie *et al. [3]*. YigI is presently annotated as “uncharacterized protein YigI” (Uniprot [16]) and as a “putative thioesterase” in the Ecocyc database [17]. Thus, YigI provided us with both an opportunity to test novel predictions from a high throughput application of our COFACTOR2/MetaGO pipeline for structure-guided functional annotation [18,19], and a starting point to investigate possible reasons for *E. coli* to encode so many thioesterases with overlapping substrate specificities.

By applying the predictions from COFACTOR2, we first experimentally show that YigI is a novel thioesterase present in *E. coli* that prefers medium-long chain acyl-CoA derivatives *in vitro*. These findings are consistent with non-peer-reviewed prior reports of similar activity in a master’s thesis [20]. We then show that YigI is involved in the thioesterase dependent β-oxidation pathway of 9-cis,11-trans-Octadecadienoic acid (conjugated linoleic acid (CLA)), along with the previously identified TesB and FadM [3]. Bioinformatic analysis further demonstrated that TesB, FadM, and YigI show largely uncorrelated regulation across environmental conditions, but that a switch between YigI and TesB expression is triggered by low pH, and that YigI is also induced by aminoglycoside treatment and by aerobic growth. We show that unlike TesB and FadM, which are broadly distributed across the tree of life, YigI appears specific to ɣ-proteobacteria, and likely plays an important and specific role under natural conditions in permitting the metabolism of unusual carbon sources.

## Methods

### COFACTOR2

The methods employed for COFACTOR2 can be found in detail in its publication [19]. Briefly, COFACTOR2 was used to assign gene ontology (GO) terms for all proteins present in the *E. coli* genome. For each predicted GO term -- molecular function (MF), cellular component (CC) and biological process (BP) -- Cscore^GO^ values are given, which corresponds to the confidence score for a particular GO term. Protein homologs are found from the predicted iTASSER structure and the Protein Data Bank (PDB) search, which classify active site residues and ligand binding sites. COFACTOR2 outputs the top 5 enzyme homologs for the query protein and gives a Cscore^EC^, which is the confidence score for the Enzyme Commission (EC) number prediction.

### Molecular Cloning and Protein purification

Gibson assembly [21] was used to transfer *yigI* from *E. coli* MG1655 into a His-SUMO-TEV (HST) plasmid backbone (Addgene product number: 48313). Each HST construct was grown in LB to an OD_600_ of 0.4 at 37°C. Cells were then cooled at 4°C for 30 minutes, then a final concentration of 0.5 mM IPTG was added to the culture to induce protein expression. Induction occurred overnight at 18°C and cell pellets were harvested the following morning following 16 hours of induction. Pellets were stored at −80°C until purification. Pellets were solubilized in lysis buffer (40 mM HEPES pH 7.3, 375 mM NaCl, 0.1% Triton X-100, 0.5 mM EDTA, 1 mM DTT, 20 mM imidazole, 1x Protease Inhibitor Cocktails (Roche)) plus 0.5 mg/mL lysozyme at 4°C, then sonicated and pelleted. The supernatant was batch bound with pre-equilibrated nickel beads for 2 hours at 4°C and then loaded onto a gravity column. The beads were washed with binding buffer (40 mM HEPES pH 7.3, 375 mM NaCl, 0.1% Triton X-100, 0.5 mM EDTA, 1 mM DTT, 10% glycerol and 20 mM imidazole), then the imidazole concentration was increased in a stepwise fashion: 50 mM, 100 mM, 150 mM, 200 mM, 250 mM, 300 mM, 350 mM, 400 mM, 500 mM, and 1 M. Protein began to elute following the addition of 150 mM imidazole. Appropriate elution fractions were then collected and concentrated in TEV cleavage buffer (50 mM Tris pH 7.9, 150 mM NaCl, and 2 mM DTT). 5U of TEV protease was added and cleavage occurred overnight at 4°C. The tag and protein of interest were separated by nickel affinity. The flow through containing the protein of interest was then concentrated in storage buffer (40 mM HEPES pH 7.3, 200 mM NaCl, 1 mM DTT, and 10% glycerol), and the final protein product was tested for activity. Following purification, protein concentration and purity were analyzed by the Bradford assay and Silver Staining, respectively. Proteins were then concentrated to approximately 50 uM and stored in 50% glycerol, aliquotted, flash frozen with liquid nitrogen, and stored at −80°C until testing.

### DTNB assay

Ellman’s Reagent, or 5,5-dithio-bis-(2-nitrobenzoicacid) (DTNB), was utilized to chromogenically detect the release of free thiols due to thioesterase activity [22]. The activity buffer contained: 20 mM HEPES pH 7.3, 50 mM NaCl, 5 mM MgCl_2_, 1 mM CaCl_2_, 0.1% Triton X-100, 5 mM MgCl_2_ and 1 mM CaCl_2_ 0,.4 mM DTNB, and varying concentrations of CoA-substrates. To determine the specificity of YigI, various CoA substrates with different carbon lengths and aromaticity were tested, including acetyl-CoA, butyl-CoA, decanoyl-CoA, benzoyl-CoA, and palmitoyl-CoA (all purchased from Sigma).

### Amidase and Lipase assays

4-nitro-7-chloro-benzofurazane (NBD-Cl) [23] was utilized to evaluate potential amidase activity for YigI. 100 nM YigI was incubated with 50 uM taurocholate and 40 mM NBD-Cl for 1 hour on a microplate reader. No amidase activity was detected (**Supplementary Data S1**). Furthermore, YigI was tested for lipase activity by using the chromogenic *p*-Nitrophenyl (pNP) assay [24]. No lipase activity was detected against pNP-acetate, -butyrate, and -palmitate (**Supplementary Data S2**).

### Activity analysis

Assays were analyzed with the GraphPad Prism 9 software under the simple linear regression feature to calculate slope values for each condition tested. Active site titration (AST) data was excluded once saturation was reached. In supplementary files, data with asterisk (*) next to a specific number were not included in the present analysis. Slope values were then plugged into the nonlinear regression (curve fit) kcat and Michaelis-Mentin enzyme kinetics feature in GraphPad. When using the kcat feature for kinetic analysis, Et values were set to input enzyme concentration.

### P1vir transduction

We obtained single gene knockouts from the Keio Collection [25] with each gene of interest replaced by a kanamycin resistance marker, and created all possible combinatorial knockouts of *yigI, tesB* and *fadM* using P1vir transduction [26] into our baseline strain (*E. coli* MG1655). The kanamycin resistance markers were then removed using the pCP20 plasmid as described by Datsenko and Wanner [27] to yield strains with in-frame deletions of the genes of interest, in the MG1655 genetic background. Strains were verified by colony PCR of the deletion locations.

### Carbon source growth assay

Knockout strains were grown as described by Nie, et al [3]. Briefly, cultures were grown in LB media at 37°C overnight, then diluted 5-fold in M9 minimal media containing 2 mM MgSO_4_, 10 μM CaCl_2_, 1 μM FeCl_3_ 1% (w/v) tryptone, 0.4% (v/v) Triton X-100, and 0.1% (v/v) oleic acid. Cell growth was measured in an Epoch 2 microplate spectrophotometer (BioTek Instruments) at 37°C with 237 cpm continuous shaking until the median optical density (OD) reached 1.00 on a 1-cm pathlength at 600 nm, which corresponds to an unblanked OD_600_ of 0.464 on the Epoch spectrophotometer. Cultures were then diluted 20-fold in M9 minimal media containing 2 mM MgSO_4_, 10 μM CaCl_2_, and 1 μM FeCl_3_ and supplemented with either 0.1% (w/v) glucose, 0.1% (v/v) oleic acid, 0.1% (w/v) palmitic acid, or 0.1% (v/v) 9(Z),11(E)-conjugated linoleic acid (CLA) before growth in the microplate spectrophotometer. Triton X-100 (0.4%, v/v) was used to solubilize oleic acid and CLA and was also added to the media containing glucose. Palmitic acid and CLA were solubilized with methyl-β-cyclodextrin (1:6 molar ratio). Cell growth occurred at 37°C with 355 cpm double orbital shaking. The OD_600_ was recorded every 10 minutes until cell growth reached stationary phase.

### Carbon source growth assay analysis

Cell growth was analyzed with a Bayesian multilevel mixed effects model using the *brms* package in R [28,29]. The *growthcurver* package was used to calculate empirical area under the growth curve (AUC) between 4 and 24 hours for each condition to quantify growth potential [30].

This time range captures logarithmic to stationary growth while minimizing initial fluctuations in OD_600_, as illustrated in **Supplementary Figure S1**. Log_2_-transformed AUC data were modeled assuming t-distributed errors. The central tendency was estimated based upon fixed effects terms for media type (*β*_*media*_) and the interaction between strain genotype and media condition (*β*_*strain:media*_), plus a random effect term that accounts for variation between experimental replicates by media condition (*γ*_*media:replicate*_). The model formula is as follows:

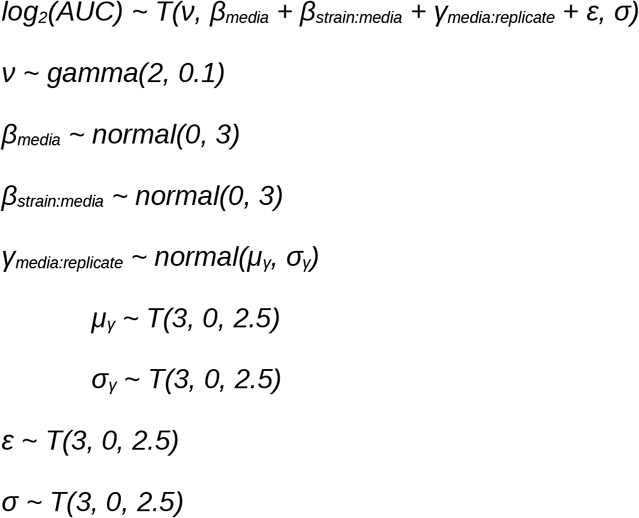

Here the order of operands to T() are the number of degrees of freedom, mean, and scale. The cumulative effect of strain genotype and media type on cell growth is of primary interest in this analysis. Based upon prior experience with *E. coli* growth rates in various carbon sources, we specified a mildly informative normal prior for the *β*_*media*_ coefficients (see above); the prior reflects our expectation that differences in growth between media types will fall within two standard deviations of the mean while still allowing for the possibility of greater differences. We used the same mildly informative prior for the *β*_*strain:media*_ coefficients based upon published data from Nie, *et al* [3]. Default priors from the *brms* package were used for all other parameters since they cover the physiologically relevant range and we have no strong reason to use different priors.

Modeling was performed using RStudio (version 1.2.5033) and Markov chain Monte Carlo (MCMC) sampling was implemented in Stan (version 2.21.0) using the *brms* package (source code given in **Supplementary Data S3**) [28,29]. As per the *brms* default specifications, MCMC sampling was performed with four chains, 2000 iterations, and 1000 burn-in steps. The model fit yielded no warning messages and ^*R* < 1.1 for all posterior parameter estimates, indicating chain convergence.

We performed pairwise comparisons for each interaction term and the corresponding WT base effect per media condition using the hypothesis function of the *brms* package [28,29]. We also performed pairwise comparisons among the interaction terms for double and triple knockout strains of a given media condition (**Figure 3D**). Comparisons were evaluated for significance based on the posterior probability of a non-zero difference in the observed direction and a difference exceeding ±log_2_(1.5). The latter threshold delineates values of practical equivalence to zero on the basis of physiologically meaningful differences.

### Transcriptome analysis

*E. coli* gene expression data at the transcript level was obtained from COLOMBOS v3.0 [31] (accessible at http://colombos.net/) as a downloaded compendium. COLOMBOS curates publicly available microarray data and homogenizes both the experimental annotations and the format for quantifying gene expression changes. The basic unit in COLOMBOS for comparing gene expression is a pair of test-versus-reference samples (“contrast”) whose log2-fold-changes were calculated.

To maximize the comparability between the COLOMBOS data and the native genetic environment of *E. coli* strain MG1655 used in our experiments, we filtered the test-versus-control pairs measured for gene expression changes (contrasts) based on their experimental annotations, to: 1) include only experiments with three K-12 strains (MG1655, BW25113, and W3110), 2) exclude experiments with genetic manipulations shared by the test and control samples (i.e. gene deletions, overexpressions, or plasmids), 3) include only experiments designed either with identical growth media annotations for the test and control samples or to compare gene expression in minimum media versus in rich media, and 4) include only experiments with none or only one varying annotation between the test and control samples (i.e. only comparisons among replicates or in single-variable designs). After the screening of experiment designs based on the strains, genotypes, consistent growth media, and well-annotated interpretable experimental designs, 1147 out of 4077 available contrasts were surveyed in our comparison of gene expressions of the thioesterases of interest. For each gene, the set of log fold changes arising from COLOMBOS from the contrasts of interests were converted into gene-wise robust z-scores *z* following the equation:

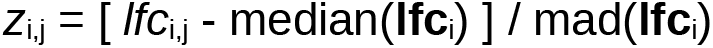

Where **lfc** denotes the log fold change matrix from COLOMBOS, *i* denotes genes, *j* denotes conditions, and mad() is the median absolute deviation.

### Gene conservation analysis

To obtain the sequence alignments and identities used in **Figure 4C**, we applied CLUSTAL OMEGA with default settings to the four thioesterase sequences under consideration. For YigI and FadM, the full amino acid sequences present in UNIPROT were considered; for TesB, we divided the protein roughly at the midpoint between the two thioesterase domains, considering residues 1-120 as the first half and the remainder as the second half. The fractions of aligned positions with identical residues were then reported. For structural alignments, we used TM-align to perform pairwise alignments of the same protein segments, using the crystal structures for TesB and FadM (PDB codes 1C8U [32] and 1NJK [33] for TesB and FadM, respectively), and our I-TASSER structural model (**Supplementary Data S4**) for YigI. As TM-align provides TM scores normalized by the lengths of both input sequences, we averaged the two provided TM scores to provide the results shown in **Fig. 4C**.

To test the conservation of our thioesterases of interest across *E. coli*, we applied a tblastn search (using default settings) to a custom database consisting of all 13,968 Genbank genomes with species name “*Escherichia coli*” for which coding sequence fasta files were available. We requied at least 80% sequence identity over a 100 amino acid or longer alignment in order to count as a hit. To search for potentially distant homologues of the *E. coli* acyl CoA thioesterases across the tree of life, we performed psiblast searches (using default parameters) against the NCBI landmark database, for each of the four protein chains defined above. We used *E. coli* O157:H7 sequences as representatives of *E. coli* for conservation analysis since the K12 sequences of our query proteins were used as inputs. The sequence identities between our query proteins and the highest-identity hit found in each database species are shown in **Fig. 4D.**

## Results

### YigI structurally resembles well-defined hot-dog fold thioesterases

YigI is a 155 amino acid, 17.163 kDa, putative thioesterase from *E. coli* strain K12. Currently, there is no published experimental information for *yigI* present on the EcoCyc database [17]. We predicted the structure of YigI using I-TASSER [34], and then ran COFACTOR2 [19] in conjunction to predict the characteristics and function of YigI.

Structurally, YigI aligned very well to a thioesterase from *Shewanella oneidensis* (TM-score = 0.95; Uniprot ID: Q8E9M7; PDB: 3E8P). Notably, YigI is predicted to possess a hot dog fold (**Fig. 1A**), which is a canonical structure for thioesterases [35]. Furthermore, the identified likely homologous protein from *Shewanella oneidensis* shares 46.7% sequence identity to YigI and 62.6% sequence similarity. GO terms were predicted for YigI (**Fig. 1B-D**). There is very high confidence that YigI localizes in the cytosol **(Fig. 1B**; Cscore^GO^ = 1.00), which corroborates previous predictions [36]. For molecular function (MF) analysis (**Fig. 1C**), YigI scored highly for catalytic activity and hydrolase activity acting on ester bonds with Cscore^GO^ values of 0.95 and 0.78, respectively. Finally, for biological process (BP) GO terms (**Fig. 1D**), YigI is strongly predicted to have cellular and metabolic roles (Cscore^GO^ = 0.94 and 0.91, respectively). Given this initial data, there is strong computational evidence that YigI is a thioesterase. We set out to purify and characterize YigI *in vitro*.

**Figure 1:**
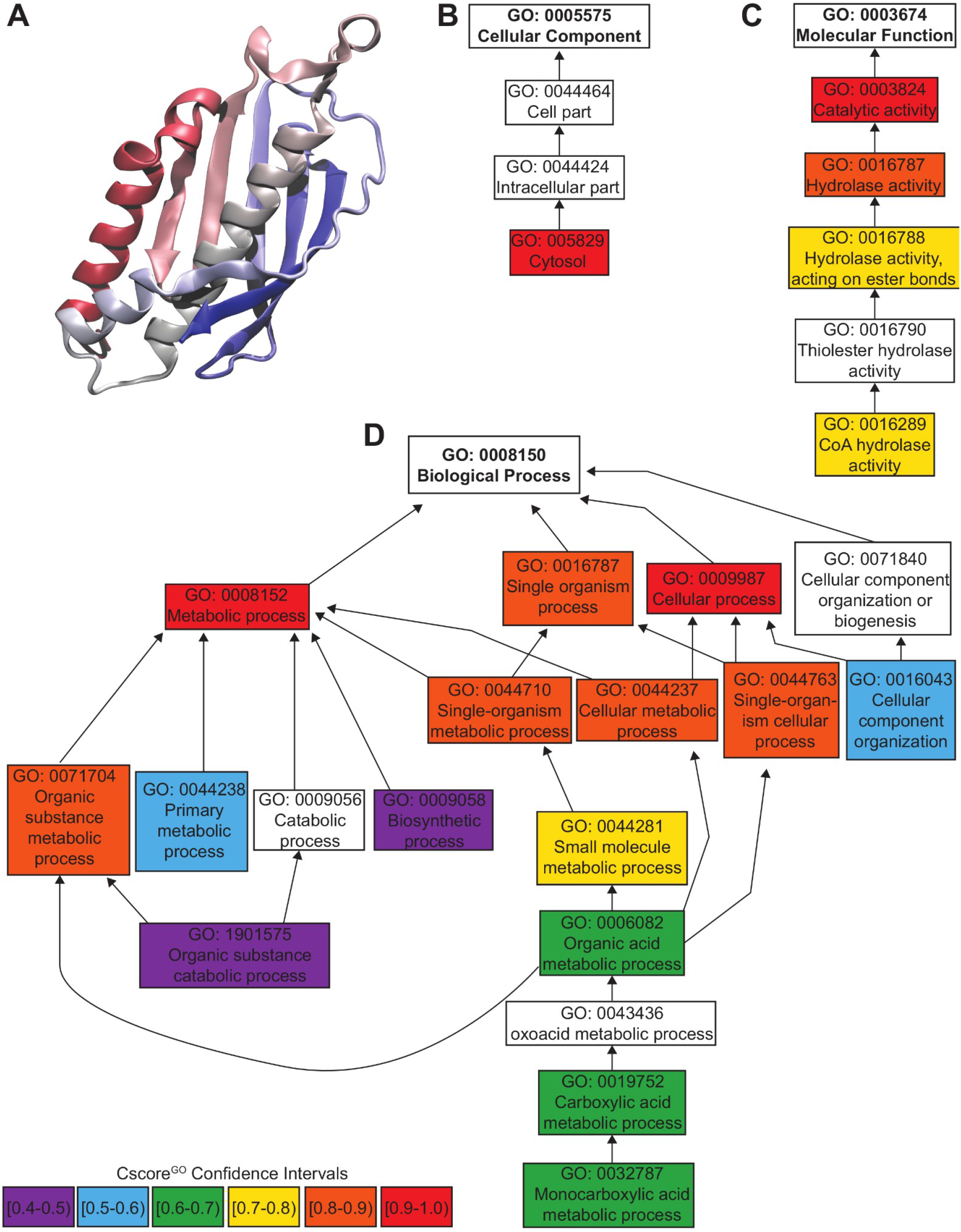
COFACTOR2 function predictions for YigI (full data available as **Supplementary Data S5**). (**A**) Predicted crystal structure of YigI from I-TASSER in COFACTOR2 (**Supplementary Data S4**). (**B**) Cellular component (CC) gene ontology (GO) term predictions for YigI. Confidence scores (Cscore) are color-coded as reflected in the bottom left; Cscores range from 0 to 1, with a higher score indicating higher confidence in a GO term annotation. (**C**) Molecular function (MF) GO term predictions for YigI. (**D**) Biological process (BP) GO term predictions for YigI.

### YigI displays thioesterase activity against medium to long chain acyl-CoA substrates

YigI was purified according to standard nickel and TEV gravity chromatography techniques (**Figure 2A**; **Supplementary Figure S2;** see **Methods**). Utilizing the Ellman’s Reagent, 5,5-dithio-bis-(2-nitrobenzoicacid) (DTNB), we determined that YigI is active against decanoyl-CoA, a 10-carbon long substrate, and that the kinetic parameters, K_M_ and k_cat_, for decanoyl-CoA fit with the canonical hotdog fold thioesterases with ~19uM and 1.4/s, respectively (**Figure 2B-E**). YigI also possessed activity towards palmitoyl-CoA, and we can conclude the enzyme appears to prefer medium- to −long chain acyl-CoA substrates *in vitro* (**Figure 2B and 2D-E;** raw data in **Supplementary Data S6** and **Supplementary Data S7**). No thioesterase activity was observed for acetyl-, butyl-, or benzoyl-CoA (**Figure 2B**). Through observing the predicted structure from I-TASSER, two catalytically dead mutants were designed, N53A and D69A (**Figure 2A;** primers used for knockouts are given in **Supplementary Data S8**). Indeed, both mutants are catalytically inactive against all substrates tested (**Figure 2C;** raw data are given in **Supplementary Data S9, Supplementary Data S10**, and **Supplementary Data S11**). In addition to our work, a previous masters’ dissertation corroborated that D69 is critical for YigI’s enzymatic activity, as D69Q and D69E mutants were catalytically dead [20]. This report also showed that Q56A produced inactive mutants, and reported similar kinetic parameters for decanoyl-CoA along with a detailed kinetic catalog for acyl-CoA substrates [20]. Additionally, we did not detect amidase and lipase activity for YigI *in vitro* (**Supplementary Data S1; Supplementary Data S2; see Methods**). Given this data, we have strong evidence that YigI is a thioesterase that acts on medium-long chain acyl-CoA substrates.

**Figure 2:**
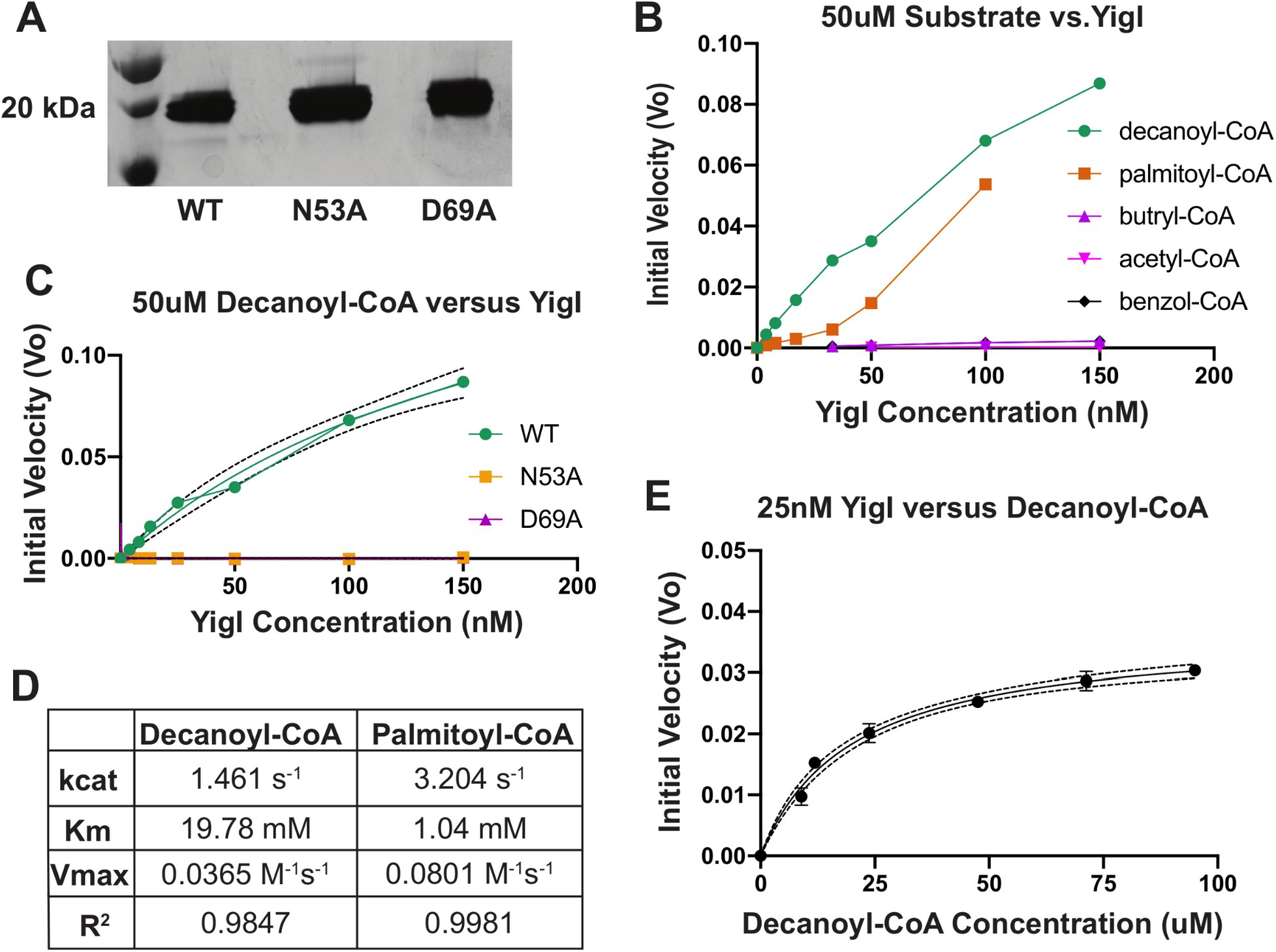
*In vitro* characterization of YigI. (**A**) Silver stain gel showing the purified YigI used in our *in vitro* experiments. WT, N53A mutant, and D69A mutants are shown, in order. (**B**) Active site titration of YigI against 50 μM of CoA substrates. Samples were run in duplicates and their averaged slopes are plotted. (**C**) Active site titration of wild type YigI, YigI-D69A, and YigI-N53A against 50μM decanoyl-CoA. Samples were run in duplicates and their averaged slopes are plotted. Dashed interval corresponds to 95% prediction intervals. (**D**) Quantification of kinetic characteristics of YigI against Decanoyl-CoA and Palmitoyl-CoA substrates. (**E**) Michaelis-Menten titration of 25nM YigI against varying decanoyl-CoA concentrations. Samples were run in triplicate, represented by error bars. Dashed interval corresponds to 95% prediction intervals. Kinetic fits arising from the data in panel E are displayed in panel D.

### Growth assays verify the role of YigI in thioesterase-dependent CLA catabolism in vivo

Following *in vitro* characterization of YigI, we sought to understand its physiological role in *E. coli* MG1655. TesB (thioesterase II) and FadM (thioesterase III) participate in CLA degradation through thioesterase-dependent β-oxidation [3], and are likely involved in the degradation of similar fatty acid substrates. When metabolized by *E. coli*, CLA undergoes three rounds of β-oxidation to 3,5-Dodecadienoyl-CoA. However, as *E. coli* lacks the dienoyl-CoA isomerase to activate the β-carbon in this fatty acyl-CoA, the pathway hits a roadblock, preventing further oxidation and potentially locking CoA into an un-metabolizable intermediate, which can be resolved by an acyl-CoA thioesterase to liberate CoA (**Figure 3A**). Previous growth experiments have demonstrated substantially reduced growth of a Δ*tesB*Δ*fadM* double knockout when CLA is the only provided carbon source, but since the double knockout was not lethal, the authors suggested the existence of at least one additional thioesterase involved in CLA catabolism [3]. Due to YigI’s *in vitro* preference for decanoyl-CoA and palmitoyl-CoA, we theorized that the enzyme might participate in this pathway alongside TesB and FadM.

**Figure 3.**
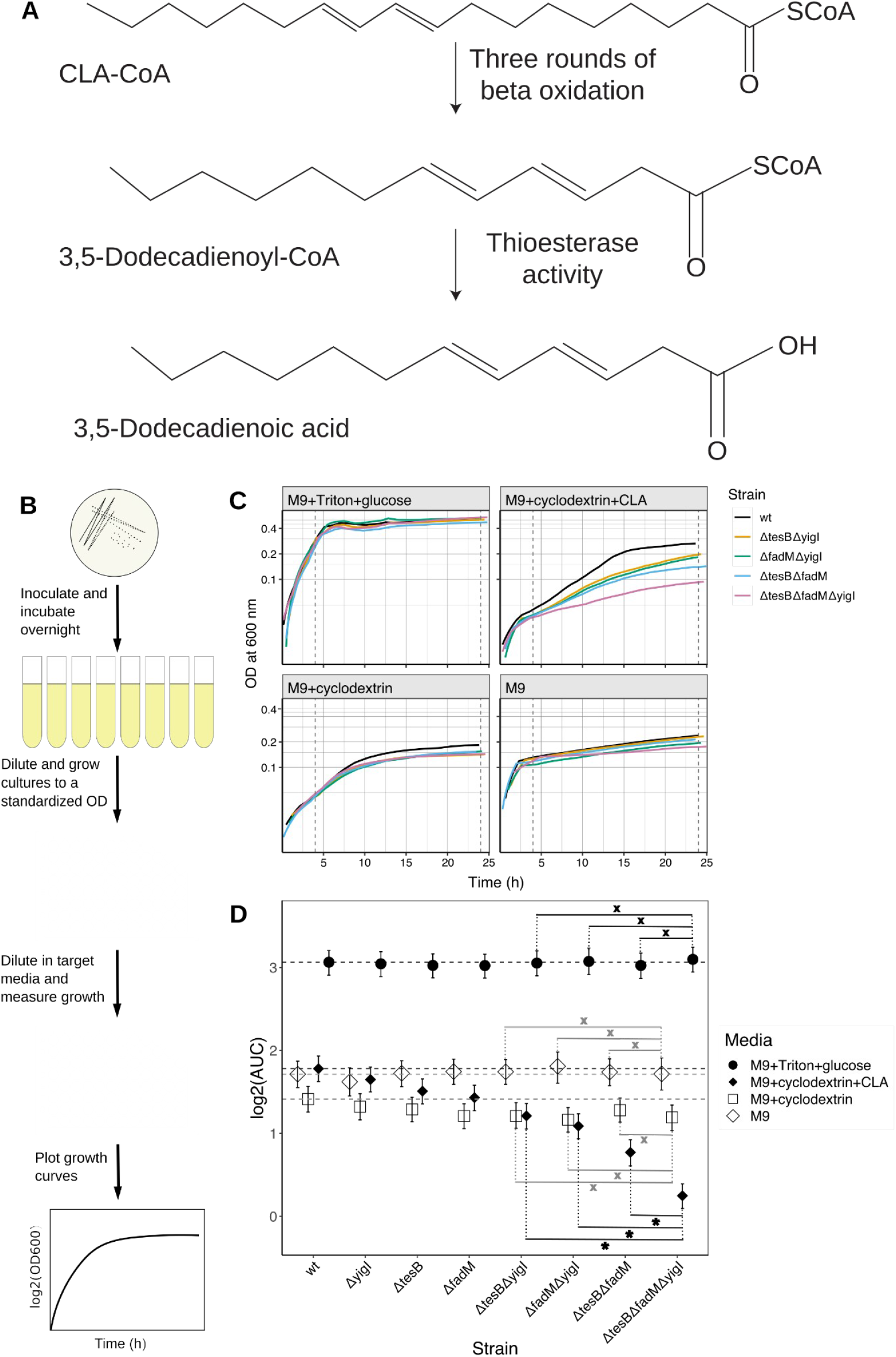
Growth of thioesterase knockout strains on CLA and other carbon sources. (A) Schematic of thioesterase-dependent metabolism of CLA, as originally described in [3]. (B) Schematic summary of the methodology used to evaluate growth phenotypes of thioesterase knockouts on different carbon sources. (C) Growth curves of the double knockouts, triple knockout, and WT strain in M9 minimal media containing glucose (0.1% w/v), CLA (0.1% v/v) and a solubilizing agent, the solubilizing agent alone, or no nutritional supplementation. (D) Conditional effects of strain genotype on log_2_-transformed area under the curve (AUC) associated with the media conditions shown in panel B. Error bars show 95% credible intervals. Dashed horizontal lines show the WT base effect for each media condition. Comparisons between the triple mutant and each double mutant for a given media condition are marked with i) an asterisk if there is a significant difference (>95% posterior probability of a difference in the indicated direction), and ii) an “x” if the difference is not biologically significant (>95% posterior probability that the absolute is less than log_2_(1.5), that is, a difference of less than 1.5-fold.

To evaluate the physiological roles of YigI, TesB, and FadM in thioesterase-dependent fatty acid catabolism, we quantified the growth of relevant knockout mutants on glucose, palmitic acid, oleic acid, or CLA as the primary carbon source *in vivo* (**Figure 3B**), as originally described in Nie et al [3]. We obtained single gene knockouts from the Keio Collection [25], and then created all possible combinatorial knockouts of Δ*yigI*, Δ*tesB* and Δ*fadM* using P1vir transduction *[26]*. Cell growth was monitored via OD_600_ measurements and plotted as growth curves. Empirical area under the curve (AUC), also known as growth potential, was used for statistical analysis.

Glucose is the preferred carbon source of *E. coli* and does not undergo fatty acid catabolism, therefore supplementation with glucose served as a positive control for bacterial growth. All strains exhibited consistent, maximal growth in glucose-containing media (**Figure 3C-D**). Palmitic acid is a saturated fatty acid and thus undergoes complete degradation via thioesterase-independent β-oxidation, and as expected, we did not observe decreased growth for any of the thioesterase knockouts in media supplemented with palmitic acid (**Supplementary Figure S1: E, I**). Approximately 10% of oleic acid is degraded via thioesterase-dependent β-oxidation, whereas the rest is degraded through the thioesterase-independent pathway [10]. Though Nie et al. reported impaired growth of Δ*tesB* and Δ*fadM* knockouts on oleic acid [3], we observed robust growth across all knockout strains (**Supplementary Figure S1: B, I**).

Unlike palmitic acid and oleic acid, partial degradation of CLA yields an acyl-CoA intermediate that cannot undergo further β-oxidation, requiring hydrolysis by a thioesterase to free the CoA molecule and excrete the unusable fatty acid [3]. Though all of the thioesterase knockouts exhibited lower growth than WT in media containing CLA, only three knockout strains had physiologically significant differences in growth relative to the WT base effect (**Supplementary Table S1, Figure 3D**). When grown in M9+cyclodextrin+CLA media, the Δ*fadM*Δ*yigI* knockout exhibited 0.6-fold change in growth potential from WT (**Table 1**, **Figure 3C-D, Supplementary Table S1**). Consistent with previous findings by Nie et al. [3], the Δ*tesB*Δ*fadM* double knockout displayed a substantial growth defect when supplemented with CLA (**Table 1**, **Figure 3C-D**).

**Table 1.** Bayesian point estimates and 95% credible intervals (CI) for interaction effects between strain genotype and media condition on log_2_-transformed area under the curve (AUC).

| Media | Strain genotype | Estimate<sup>a</sup> | Lower CI | Upper CI |
| --- | --- | --- | --- | --- |
| M9+Triton+glucose | *ΔyigI* | -0.02 | -0.13 | 0.09 |
|  | *ΔtesB* | -0.04 | -0.16 | 0.08 |
|  | *ΔfadM* | -0.04 | -0.15 | 0.07 |
|  | *ΔtesBΔyigI* | -0.01 | -0.12 | 0.10 |
|  | *ΔfadMΔyigI* | 0.01 | -0.11 | 0.14 |
|  | *ΔtesBΔfadM* | -0.04 | -0.16 | 0.08 |
|  | *ΔtesBΔfadMΔyigI* | 0.04 | -0.08 | 0.15 |
| M9+cyclodextrin+CLA | *ΔyigI* | -0.13 | -0.26 | -0.01 |
|  | *ΔtesB* | -0.27 | -0.40 | -0.16 |
|  | *ΔfadM* | -0.35 | -0.48 | -0.24 |
|  | *ΔtesBΔyigI* | -0.57 | -0.70 | -0.45 |
|  | *ΔfadMΔyigI* | -0.70 | -0.82 | -0.58 |
|  | *ΔtesBΔfadM* | -1.02 | -1.15 | -0.88 |
|  | *ΔtesBΔfadMΔyigI* | -1.54 | -1.66 | -1.41 |
| M9+cyclodextrin | *ΔyigI* | -0.09 | -0.22 | 0.04 |
|  | *ΔtesB* | -0.13 | -0.25 | 0.00 |
|  | *ΔfadM* | -0.21 | -0.33 | -0.07 |
|  | *ΔtesBΔyigI* | -0.21 | -0.33 | -0.06 |
|  | *ΔfadMΔyigI* | -0.25 | -0.37 | -0.12 |
|  | *ΔtesBΔfadM* | -0.14 | -0.26 | 0.00 |
|  | *ΔtesBΔfadMΔyigI* | -0.22 | -0.36 | -0.09 |
| M9 | *ΔyigI* | -0.09 | -0.24 | 0.06 |
|  | *ΔtesB* | 0.01 | -0.12 | 0.14 |
|  | *ΔfadM* | 0.03 | -0.09 | 0.16 |
|  | *ΔtesBΔyigI* | 0.03 | -0.1 | 0.16 |
|  | *ΔfadMΔyigI* | 0.09 | -0.08 | 0.24 |
|  | *ΔtesBΔfadM* | 0.03 | -0.11 | 0.17 |
|  | *ΔtesBΔfadMΔyigI* | 0.00 | -0.18 | 0.17 |

<sup>a</sup> Interaction coefficients are given as deviations from the WT base effect per media condition, therefore these numbers do not correspond exactly to the graphical depiction in Figure 3D.

Loss of both thioesterases known to participate in CLA catabolism yielded 0.5-fold change in growth potential (**Supplementary Table S1**). Strikingly, the Δ*tesB*Δ*fadM*Δ*yigI* mutant exhibited the lowest growth of any knockout strain on CLA, demonstrating that YigI does partially complement TesB and FadM function in thioesterase-dependent β-oxidation *in vivo* (**Table 1**, **Figure 3C-D**). The triple mutant was associated with a 0.3-fold change in growth potential relative to WT (**Supplementary Table S1**). Additionally, the growth exhibited by the triple mutant was significantly lower than that of any of the double knockouts in M9+cyclodextrin+CLA media (**Supplementary Table S1, Figure 3D**).

It is important to note that we observed bacterial growth even in unsupplemented M9 minimal media (**Figure 3C-D**). We believe that the cells likely utilized residual amounts of tryptone transferred from the pre-growth media to the target media via the inoculum for nutrition. We also observed low growth in media with cyclodextrin alone (**Figure 3C-D**), therefore we attribute growth to residual nutrients present in the back dilution rather than utilization of cyclodextrin itself as a carbon source. Interestingly, despite evidence indicating that residual nutrients are present in target media, the Δ*tesB*Δ*fadM* and Δ*tesB*Δ*fadM*Δ*yigI* knockouts exhibit lower growth in CLA-containing media than in unsupplemented media (**Figure 3C-D**). Thus, the apparent toxicity of blocked CLA catabolism is best explained by depletion of intracellular CoA and/or the presence of the non-metabolizable product, rather than the cells’ inability to utilize CLA as a carbon source. We also note that aside from the cases discussed above, several other condition-genotype interactions showed very small but statistically significant effects, as detailed in **Supplementary Text S1**, but the magnitude of these effects was too small to appear biologically noteworthy.

### YigI expression is induced by low pH and by aminoglycoside treatment

In order to obtain additional insight into the functional distinctions between *E. coli* thioesterases, and in particular into the biological role of YigI, we considered the distributions of expression levels (at the transcript level) of TesB, FadM, and YigI across all datasets present in the COLOMBOS database [31]. As shown in **Fig. 4A-B**, we observed that expression of the three thioesterases was in general not well correlated (pairwise Spearman correlations below 0.2 in all cases), and conditions that strongly induce or reduce the expression of one particular thioesterase in general do not show strong impacts on the expression of the others (with one exception that will be discussed below). To further explore the conditions that do strongly affect expression of the thioesterases of interest, we tabulated the conditions showing the largest impacts on thioesterase expression in **Table 2**. While in general the conditions with strong impacts on thioesterase expression appear unrelated across the three transcripts considered, there is a notable overlap in that low pH conditions (pH 5.5) appear to trigger both substantial increases in YigI expression and substantial decreases in TesB expression, corresponding to the cluster of points at the extreme lower-right of the density plot in **Fig. 4A**. In addition to low pH, YigI expression appears to also be increased by treatment with aminoglycosides, and based on data from three different studies (NCBI GEO accession numbers: GSE21839 [37], GSE13982 [38], GSE1121 [39]), decreased under anaerobic conditions. Simply on the basis of the expression data considered here, it is not immediately clear whether the induction/inhibition of YigI under some conditions (and in particular, the tradeoff favoring YigI relative to TesB under acidic conditions) reflects the efficacy of the enzymes themselves under those conditions, the accumulation of a substrate upon which YigI is particularly effective under those conditions, or possibly even an ecological link in which a favored substrate for YigI is often encountered under those conditions in the native ecology of *E. coli* (in which case low pH or aminoglycosides might serve as a cue for anticipatory regulation [40]).

**Table 2:** Conditions showing the largest log fold-changes in expression of YigI, TesB, and FadM across the COLOMBOS database. In each case we calculated robust z-scores for the observed log fold changes for each gene of interest, and then considered the twenty contrasts with the most positive and most negative z-scores for each transcript. After excluding all contrasts for which one or both strains under consideration was evolved or genetically modified, we manually consolidated the remaining top/bottom 20 cases under a human-readable description of the key intervention present in those samples (“Condition”). For each such condition, we give the largest magnitude robust z-score observed in the “Max |rz|” column, and the number of instances of a similar condition in the top/bottom 20 contrast set for that transcript in the “# in top/bottom 20”. Note that the total condition count in that column may not sum to 20 due to exclusion of genetically manipulated strains. A complete enumeration of the COLOMBOS contrasts present in these top/bottom 20 lists are provided in **Supplementary Data S12**.

| Condition | Max \|rz\| | # in top/bottom 20 |
| --- | --- | --- |
| Highest expression of YigI |  |  |
| Gentamicin | 11.3145438 | 3 |
| pH 5.5 | 8.654750855 | 12 |
| Aerobic growth (vs anaerobic growth) | 7.817330052 | 1 |
| Kanamycin | 7.447106537 | 3 |
| Lowest expression of YigI |  |  |
| Polyhexamethylene biguanide | -14.863912 | 1 |
| Anaerobic growth (vs aerobic growth) | -11.37200684 | 7 |
| Carbon monoxide | -8.177883356 | 2 |
| Highest expression of TesB |  |  |
| Carbon monoxide | 7.408683056 | 1 |
| H2O2 | 7.101972983 | 3 |
| Glucose → lactose diauxie | 6.797398774 | 2 |
| Bacillus VOCs | 6.746138037 | 3 |
| High salt | 6.509875167 | 3 |
| Stationary phase | 5.847353643 | 3 |
| Sodium nitrate (vs oxygen) | 5.319684166 | 1 |
| Gentamicin | 5.181404057 | 1 |
| Lowest expression of TesB |  |  |
| PH 5.5 | -20.72534497 | 18 |
| Glucose → acetate diauxie | -13.73765225 | 1 |
| Biofilm | -13.58024761 | 1 |
| Highest expression of FadM |  |  |
| Stationary phase | 22.42443649 | 8 |
| Acetate (vs glucose) | 13.47665126 | 2 |
| High salt | 12.33866643 | 3 |
| Single amino acid carbon sources (vs glucose) | 10.56109548 | 4 |
| Glass wool | 10.36994691 | 1 |
| Glucose → lactose diauxie | 9.54581259 | 1 |
| Lowest expression of FadM |  |  |
| Rapid growth | -22.97732819 | 8 |
| S-nitrosoglutathione | -9.375090717 | 1 |
| Carbon starvation | -6.184400504 | 1 |
| Bacillus VOCs | -5.983231688 | 1 |
| Isobutanol | -5.877978805 | 1 |

**Figure 4:**
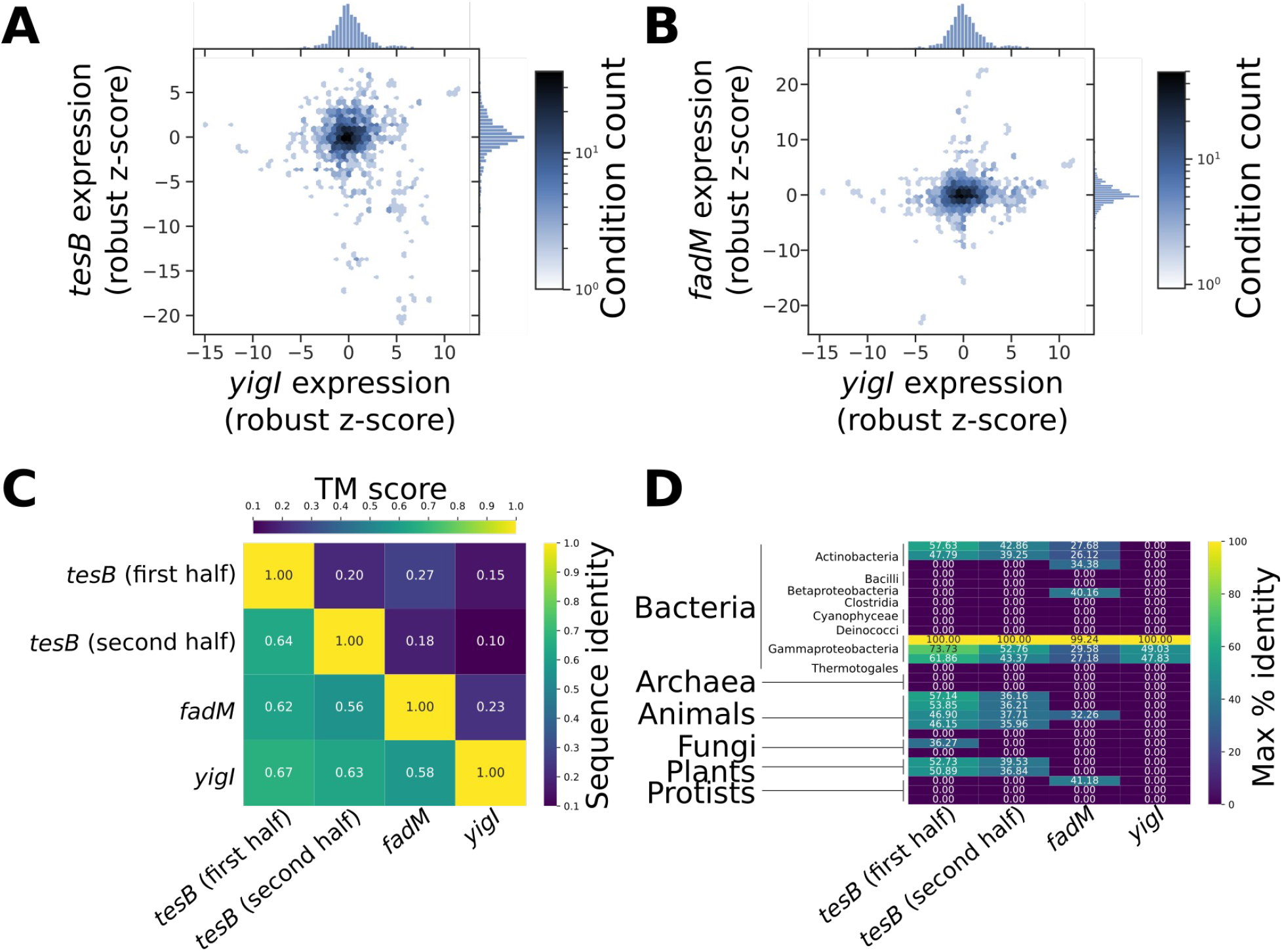
Bioinformatic analysis of the expression and conservation of *E. coli* acyl-CoA thioesterases. **(A)** Hexbin density plot showing the relative distributions of expression changes for *yigI* and *tesB* in response to the range of conditions represented in the COLOMBOS database. **(B)** As in **A**, comparing *yigI* and *fadM* expression. **(C)** Structural (lower triangle) and sequence (upper triangle) similarities between the four *E. coli* thioesterase domains considered here; TM scores are averaged across normalizations by the longer and shorter protein in each case. **(D)** Sequence identities of the best hit (identified via psiblast) in the landmark database for each of the *E. coli* MG1655 thioesterase domains considered here. “0” is shown in cases where no detectable homologs were present passing our E-value threshold.

### YigI is a gammaproteobacteria-specific thioesterase

To place our findings regarding YigI in the broader biological and evolutionary context of acyl-CoA thioesterases, we searched for identifiable homologs of *yigI* and similar thioesterases across both *Escherichia coli* and a broader sampling of organisms. As TesB contains two structurally similar thioesterase domains [32] we considered separately the N-terminal and C-terminal domains of TesB (which we refer to below as “first half” and “second half”, respectively). Beginning by simply considering the similarities between the various acyl CoA thioesterases within *E. coli*, we used both sequence alignments (using CLUSTAL OMEGA [41]) and structural alignments (using TM-align [42]) to compare the *E. coli* MG1655 versions of the proteins. We found that while all four thioesterase domains had substantial divergences in sequence, their folds were remarkably similar, highlighting the conservation of the ‘hot dog’ thioesterase fold and functional features of the proteins despite the large divergence in primary sequence (**Fig. 4C**). Applying a tblastn [43] search across a set of 13,953 *E. coli* genomic sequences available from NCBI, 13,798 (98.8%) of them contained identifiable copies of *yigI*, *fadM*, and both halves of *tesB* (in all cases requiring >80% amino acid sequence identity to the MG1655 versions of each protein over a 100 amino acid or longer stretch). Thus, the presence of all three of *yigI*, *fadM*, and *tesB* appears ubiquitous within *Escherichia coli*. Looking more broadly, we applied psiblast [44] searches against the landmark database of representative species to track the conservation of the *E. coli* acyl CoA thioesterases across the tree of life. As shown in **Figure 4D**, whereas both *tesB* and *fadM* have identifiable homologs across several kingdoms, *yigI* appears only amongst the gammaproteobacteria; of the 29 species included in the database, only three (*E. coli*, *P. aeruginosa*, and *S. oneidensis*) has identifiable *yigI* genes. Thus, *yigI* appears to be taxonomically restricted to one particular bacterial class, but is apparently widespread within that class, and likely plays an important role in the varied metabolisms of gammaproteobacteria.

## Discussion

Here, we complete a missing gap in understanding the physiological role of *E. coli* thioesterases by defining at least one role of the poorly characterized acyl-CoA thioesterase YigI. Guided by computational structure/function predictions, we showed that YigI is indeed functional as an acyl CoA thioesterase, while lacking general lipase or amidase activity. COFACTOR2 predictions guided our *in vitro* characterization, especially via site-directed mutagenesis in which we created two catalytically inactive mutants (N53A and D69A). From a physiological perspective, we find that as previously speculated [3], *E. coli* in fact possesses at least three distinct acyl-CoA thioesterases with overlapping substrate specificities, all of which are capable of contributing to salvage growth in the presence of CLA. YigI joins the company of TesB and FadM in providing the apparently overlapping functionality required to permit growth in the presence of CLA.

In our growth studies including CLA, it is notable that although growth is arrested, the triple knockout (Δ*yigI*, Δ*tesB* and Δ*fadM*) is not lethal when grown on CLA; it shows severely limited, but not completely halted, growth. One possible explanation for this observation is that there may be another thioesterase that can metabolize CLA in a very limited manner (presumably by acting on the 3,5-dodecadienoyl-CoA intermediate shown in **Fig. 3A**). One could speculate that this thioesterase may be YciA, as this thioesterase harbors broad substrate specificity, and future experiments could evaluate the competency of the quadruple knockout of Δ*yigI*, Δ*tesB*, Δ*fadM*, and Δ*yciA* in CLA catabolism. Another possibility for the very slow residual growth of the triple mutant in the presence of CLA is the possibility that *de novo* CoA synthesis by the cells provides a constant flux of fresh cofactor, which can then be used by the cells to support growth even though it is eventually removed circulation when it becomes trapped in 3,5-dodecadienoyl-CoA – in this scenario, CoA synthesis itself becomes strictly growth limiting. The possibilities outlined here could be distinguished in the future through characterization of the growth properties of a *yigI/tesB/fadM/yciA* mutant in the presence of CLA, compared with genetic or environmental interventions increasing CoA synthesis (as exogenous CoA cannot be directly taken up and used by *E. coli* [45]).

The observed compensatory nature of the three acyl CoA thioesterases that we considered (YigI, FadM, and TesB) suggests that there is mechanistic overlap between the enzymes, and that on top of CLA degradation, it is likely YigI participates in the additional pathways alongside TesB and/or FadM due to their relatively broad substrate affinities. The notion that *E. coli* harbors similar thioesterases that are capable of catalyzing overlapping reactions highlights the importance these proteins have in *E. coli* physiology: if one thioesterase is incapable of fulfilling its function, another can faithfully replace it. Examination of the condition-dependent expression profiles of YigI, TesB, and FadM demonstrated that their expression is mostly uncorrelated, with the exception that acidic conditions (extracellular pH 5.5) appear to trigger a switch to high YigI expression and low TesB expression. At present, the reasons for *E. coli* to bear three apparently functionally similar acyl CoA thioesterases remains unknown; the most likely explanation seems to us to be the possibility that the distinct substrate preferences of the enzymes make each thioesterase more or less suited for the set of potential substrates that cells might encounter under different conditions. Future investigations more specifically characterizing the activity profiles of TesB, FadM, and YigI against different thioester substrates might provide further insight into the reactions most readily catalyzed by each, and the reason for their differing condition-dependent expression profiles.

Acyl-CoA thioesterases play an important physiological role in lipid metabolism in all organisms. We found by sequence analysis that while *E. coli* TesB and FadM have identifiable homologues throughout the tree of life, YigI appears to be mostly restricted to ɣ-proteobacteria (but is broadly distributed within that class). The particular benefit conveyed by YigI will likely become clearer once the differences in its substrate specificities with those of TesB and FadM are more fully understood; one likely possibility is that YigI provides robust catalysis of some substrate that is not acted upon efficiently by TesB and FadM. The presence of an apparently narrowly distributed acyl CoA thioesterase in the ɣ-proteobacteria also raises the likely possibility that similar specialized thioesterases are common in other bacterial groups, and might play important roles in bacterial survival in the presence of a wide range of potential carbon sources and environmental hazards.

More generally, our findings illustrate the utility of computational function predictions, particularly those informed by and providing structural insights, in understanding the physiological roles of poorly annotated proteins. The information gap between sequence and function for unknown proteins will continue to grow as researchers sequence more genomes without understanding the activity of the encoded proteins. Ultimately, it is not feasible to experimentally characterize all putative proteins in every organism, highlighting the importance of ongoing efforts to improve and evaluate computational function prediction methods [46]. Comprehensive examinations of the functionalities encoded by organisms of interest can both reveal new proteins of biotechnological utility, and pinpoint metabolic weak points which might be targets for future antimicrobial therapies.

In summary, YigI is a cytosolic thioesterase in *E. coli* MG1655 that favors longer chain acyl-CoA derivatives and participates in the degradation of conjugated fatty acids such as CLA, alongside the other cytosolic thioesterases TesB and FadM. Further investigation is likely needed to determine the extent to which these three thioesterases may each be specialized toward different substrates encountered by *E. coli* in their environment, or to otherwise unravel the reasons underpinning the overlapping substrate specificities but distinct regulatory profiles of YigI, TesB, and FadM.

## Acknowledgments

We are grateful to Rachel Nicholas for testing a range of other carbon sources for the effects of *yigI* knockouts, to Benjamin Ide for assistance with strain construction/maintenance, and to members of the Freddolino lab for many helpful discussions. This work was supported by NIH grant R01 AI13467801. TP was additionally supported by NIH R25 GM086262.

## Supplementary Material

### Supplementary Data

**Supplementary Data S1**: Raw data values and experimental setup for NBD-Cl amidase assay for YigI.

**Supplementary Data S2**: Raw data values and experimental setup for pNP-lipase assay for YigI.

**Supplementary Data S3**: R code for the Bayesian multilevel mixed effects model used to analyze the carbon source growth assay data.

**Supplementary Data S4**: I-TASSER model of YigI, based on the MG1655 protein coding sequence.

**Supplementary Data S5**: Complete COFACTOR2 results for YigI functional annotations and binding sites, based on the I-TASSER predicted structure.

**Supplementary Data S6**: Raw data values for YigI against Decanoyl-CoA for figure 2D and 2E.

**Supplementary Data S7**: Raw data values for YigI against Palmitoyl-CoA for figure 2D and 2E.

**Supplementary Data S8**: YigI primers ordered from IDT and used for site directed mutagenesis.

**Supplementary Data S9**: Raw data values for YigI-N53A mutant titrated against 50uM decanoyl-CoA. Slope values are plotted in Figure 2C.

**Supplementary Data S10**: Raw data values for YigI-D69A mutant titrated against 50uM decanoyl-CoA. Slope values are plotted in Figure 2C.

**Supplementary Data S11**: Raw data values for YigI mutant titrated against 50uM decanoyl-CoA. Slope values are plotted in Figure 2C.

**Supplementary Data S12**: COLOMBOS contrasts for the most positive and most negative 20 robust Z scores for each of *yigI*, *tesB*, and *fadM*; these data underpin **Table 2** in the main text. The first tab of the spreadsheet gives a summary of the information contained in each column.

### Supplementary Figures

**Supplementary Figure S1**: Full graphical summary of the carbon source growth assay. (A-H) Growth curves for all thioesterase knockouts in each media condition tested. Assays were performed in triplicate. Knockouts were grown in M9 minimal media containing (A) glucose (0.1% w/v) and Triton X-100; (B) oleic acid (0.1% v/v) and Triton X-100; (C) CLA (0.1% v/v) and Triton X-100; (D) Triton X-100 alone; (E) palmitic acid (0.1% w/v) and cyclodextrin; (F) CLA (0.1% v/v) and cyclodextrin; (G) cyclodextrin alone; or (H) no supplemented carbon source. (I) Conditional effects of strain genotype on log_2_-transformed area under the curve (AUC) by media condition. Error bars show 95% credible intervals.

**Supplementary Figure S2**: Full gel from which Figure 2A is excerpted. Silver stain gel showing the purified YigI used in our *in vitro* experiments. Lanes from left to right: (1) ladder, (2) WT YigI, (3) blank, (4) YigI N53A mutant, (5) blank, (6) YigI D69A mutant, (7) blank, (8) YigI from a previous batch (not used in the present work), (9) YigI/chitin binding domain-chitin complex (not used in the present work), (10) YigI from a previous batch (not used in the present work).

### Supplementary Tables

**Supplementary Table S1:**
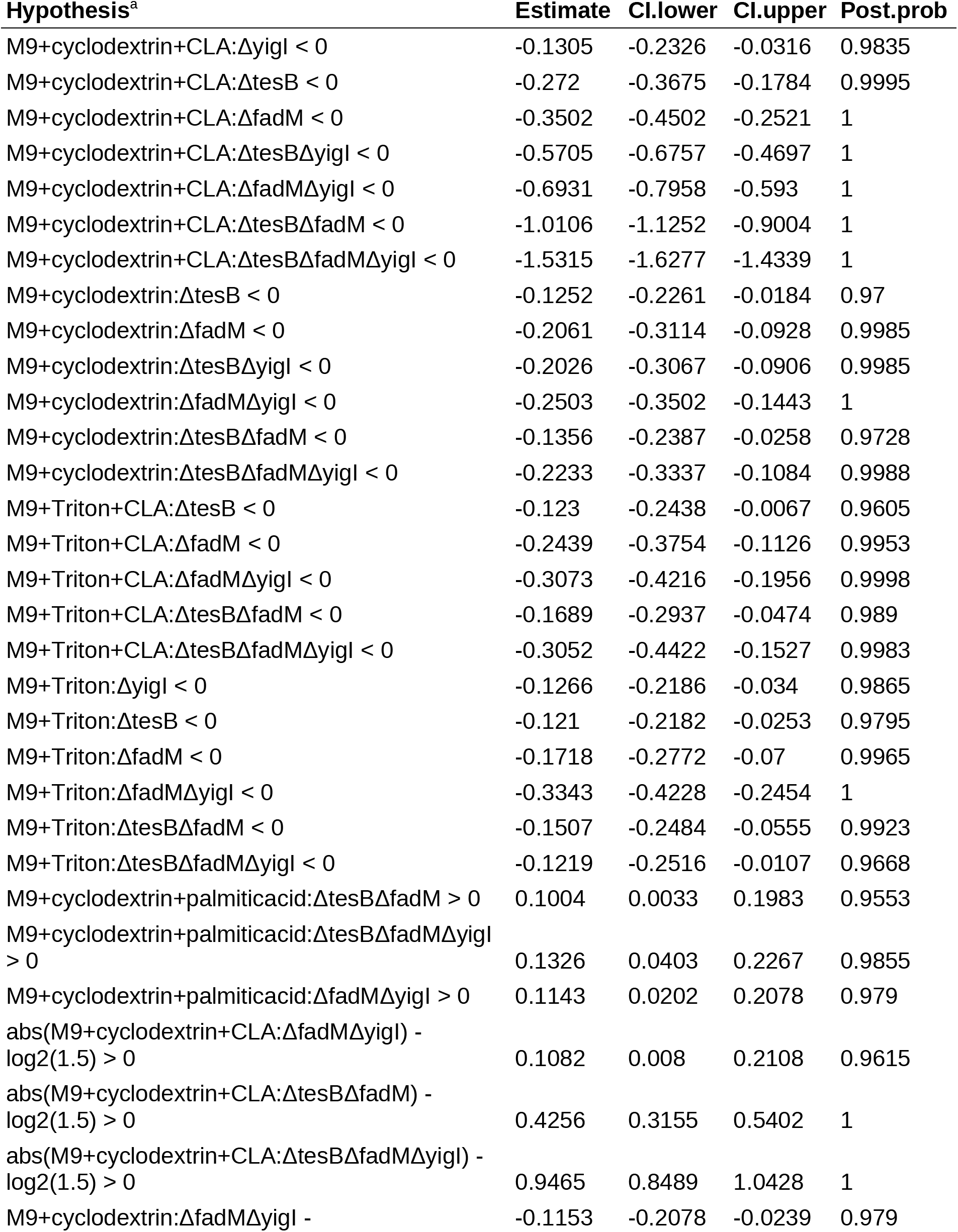
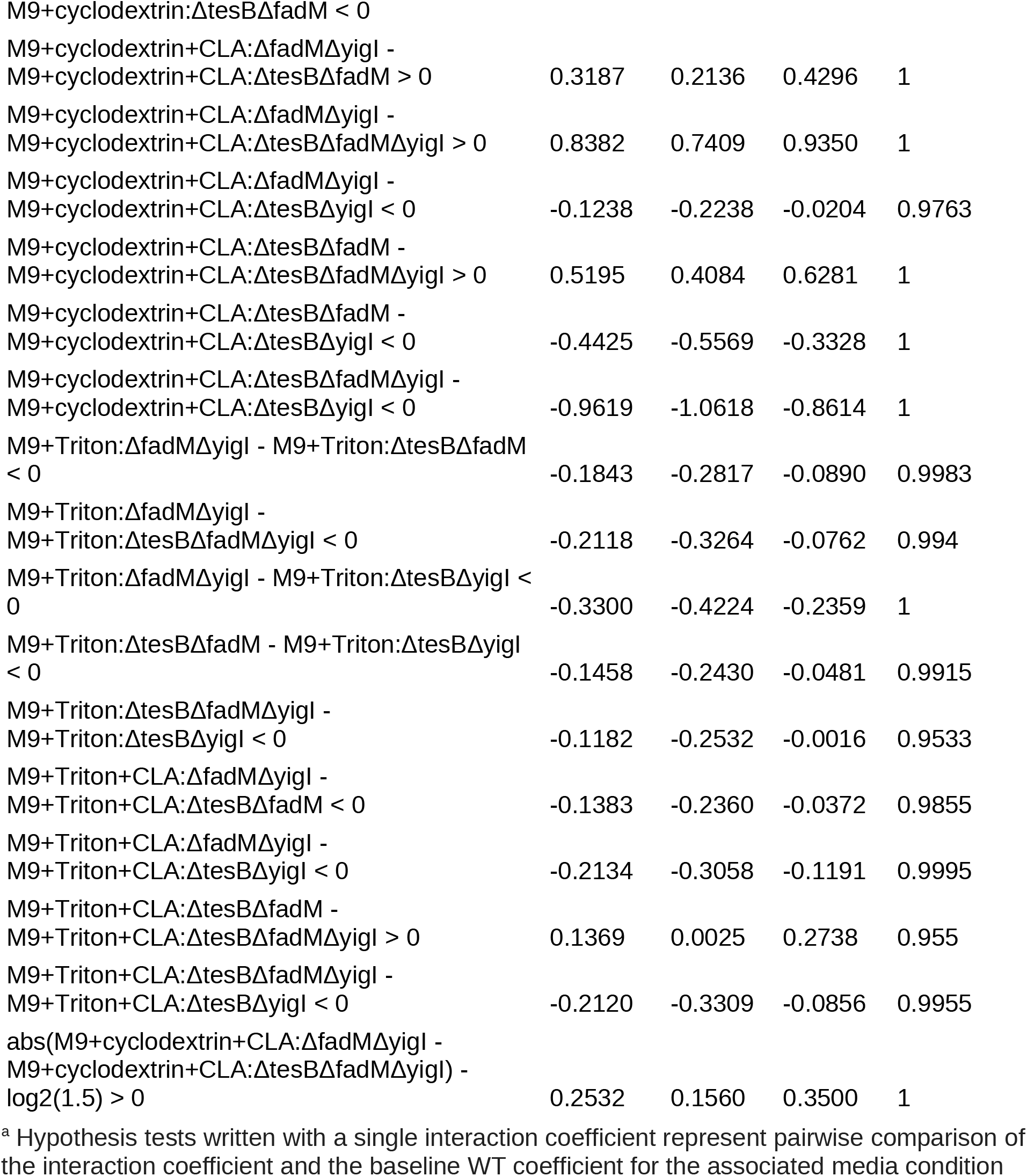
Notable pairwise comparisons between interaction terms in the form of Bayesian hypothesis tests. All interaction terms are given using the notation *media*:*genotype*, reflecting the formulation of the model itself.

### Supplementary Text

#### Supplementary Text S1

In addition to the large genotype-condition interactions between thioesterase deletions and M9+cyclodextrin+CLA media, we observed several other conditions where small changes in growth were present in some genotypes. The Δ*fadM*Δ*yigI*, Δ*tesB*Δ*fadM*, and Δ*tesB*Δ*fadM*Δ*yigI* mutants all had increased growth potential relative to WT in the presence of palmitic acid, however the differences are too small to be physiologically meaningful (**Supplementary Table S1**). In addition, the majority of knockout strains had statistically significant decreases in growth potential relative to WT in media containing CLA and Triton X-100 (**Supplementary Table S1**), however the differences are not physiologically significant nor graphically apparent (**Supplementary Figure S1: C, I**). We found similarly negligible decreases in growth for most knockouts in media containing Triton alone (**Supplementary Table S1, Supplementary Figure S1: D, I**). It is possible that Triton provides an alternative carbon source to CLA with little to no dependence on thioesterases, thus avoiding the substantial growth defects observed for thioesterase knockouts in M9+cyclodextrin+CLA media. The growth supported by M9+Triton media resembles the observed growth in unsupplemented media (**Supplementary Figure S1: I**), however, suggesting that Triton simply prevents CLA from entering the metabolic pathway that leads to poisoning of the cells.

